# Single cell transcriptomics of Atlantic salmon (*Salmo salar* L.) liver reveals cellular heterogeneity and immunological responses to challenge by *Aeromonas salmonicida*

**DOI:** 10.1101/2022.07.12.498799

**Authors:** Richard S. Taylor, Rose Ruiz Daniels, Ross Dobie, Shahmir Naseer, Thomas C. Clark, Neil C. Henderson, Pierre Boudinot, Samuel A.M. Martin, Daniel J. Macqueen

## Abstract

The liver is a multitasking organ with essential functions for vertebrate health spanning metabolism and immunity. In contrast to mammals, our understanding of liver cellular heterogeneity and its role in regulating immunological status remains poorly defined in fishes. Addressing this knowledge gap, we generated a transcriptomic atlas of 47,432 nuclei isolated from the liver of Atlantic salmon (*Salmo salar* L.) contrasting control fish with those challenged with a pathogenic strain of *Aeromonas salmonicida*, a problematic bacterial pathogen in global aquaculture. We identified the major liver cell types and their sub-populations, revealing poor conservation of many hepatic cell marker genes utilized in mammals, while identifying novel heterogeneity within the hepatocyte, lymphoid, and myeloid lineages. This included polyploid hepatocytes, multiple T cell populations including γδ T cells, and candidate populations of monocytes/macrophages/dendritic cells. A dominant hepatocyte population radically remodeled its transcriptome following infection to activate the acute phase response and other defense functions, while repressing routine functions such as metabolism. These defense-specialized hepatocytes showed strong activation of genes controlling protein synthesis and secretion, presumably to support the release of acute phase proteins into circulation. The infection response further involved up-regulation of numerous genes in an immune-cell specific manner, reflecting functions in pathogen recognition and killing, antigen presentation, phagocytosis, regulation of inflammation, B cell differentiation and T cell activation. Overall, this study greatly enhances our understanding of the multifaceted role played by liver cells in immune defense and metabolic remodeling following infection and provides many novel cell-specific marker genes to empower future studies of this organ in fishes.

## Introduction

The vertebrate liver is a multitasking organ with diverse physiological functions, including nutrient metabolism, transport and storage, growth signaling, endocrine regulation, and immunity (1). In mammals, these roles are performed by the cooperative actions of several distinct cell types including hepatocytes, cholangiocytes (epithelial cells of the bile duct), stellate cells, kupffer cells (resident liver macrophages) and lymphocytes (1). Recent advances in single cell transcriptomics have revealed functional heterogeneity within the major hepatic cell types of mammals (2-3), providing insights into liver spatial organization (4-5) while revealing cellular and molecular drivers of disease and malignancy states (6-7).

The adult liver of all vertebrates contains both immune and non-immune cells with important immunological functions (2, 7, 8-9) that support immune homeostasis and tolerance (10) and the generation of inflammatory responses upon pathogen challenge, leading to secretion of acute phase proteins (APPs) into circulation by hepatocytes (7, 11). The liver is the major site of haematopoiesis in the mammalian fetus and hence an important organ for early immune cell development (12), though this feature is not conserved in fishes (13). The multifaceted functions of the liver must demand tight coordination of different cell types to achieve appropriate responses to prevailing physiological and environmental conditions, inclusive of immune system status following pathogen challenge. Immunological functions may need to be prioritized at the cost of investment into metabolic functions in such scenarios (14). In this regard, the role of the liver in co-regulating metabolism and immunity makes it an interesting organ to understand the coordinated responses of different cell types following pathogenic challenge.

Functional genomic studies of the liver in commercially important fishes have to date applied bulk RNA-Seq and proteomics. For example, in Atlantic salmon (*Salmo salar* L.), which is among the most important aquaculture species globally (15), such work has uncovered the role of this organ in innate immune defense and the acute phase response (11,16). In the biomedical model zebrafish (*Danio rerio*), single cell transcriptomics has recently been used to reveal hepatic cellular heterogeneity, including within the hepatocyte, myeloid and lymphoid lineages (17-18). However, zebrafish are distantly related to salmonids, leaving a gap in knowledge on the role of liver cellular heterogeneity in this key group of fishes, where such information can be applied to understand and manipulate health and immune traits relevant to sustainable aquaculture and food production.

The aim of this study was to reveal the major cell lineages within the liver of Atlantic salmon and uncover the role of hepatic cellular heterogeneity in the host response to bacterial infection. Using single-nuclei RNA-Seq (snRNA-Seq), we report a comprehensive single cell transcriptomic atlas of the Atlantic salmon liver, identifying novel heterogeneity across multiple hepatic cell types. By comparing cell-specific responses in control animals to those challenged with *Aeromonas salmonicida*, we uncover a dramatic transcriptomic remodeling of hepatocytes that underpins the acute phase response, alongside major changes in gene expression specific to distinct immune cell populations.

## Materials and Methods

### Disease challenge and sampling

Animal work was carried out in compliance with the Animals (Scientific Procedures) Act 1986 under Home Office license PFF8CC5BE and was approved by the ethics committee of the University of Aberdeen. Atlantic salmon were kept in 250 L freshwater tanks at aquarium facilities of the University of Aberdeen. Water temperature was maintained at 14°C and fish were fed a commercial pellet diet at 2% body weight per day. After two weeks, twenty fish were anaesthetized using 2-phenoxyethanol (2.5ml in 10L water/ 0.0025% v/v) and given an intraperitoneal injection of either PBS (0.5 mL) (n=10), or the pathogenic Hooke strain of *A. salmonicida* (2 × 10^5^ colony-forming units / mL in PBS; 0.5 mL/fish) (n=10). Sampling was performed 24 h post-injection (after ref 11). The fish were killed using a Schedule 1 method following overdose anaesthetization using 2-phenoxyethanol (0.1% v/v) and destruction of the brain. Fish were immediately sampled, with liver samples (approx. 100mg) flash frozen on dry ice before storage at -80°C prior to snRNA-Seq library construction. Separate liver samples were placed in 1.5ml of Tri Reagent (Sigma-Aldrich) and used for quantitative PCR (qPCR) to validate the expected response in infected fish.

### Validation of immune response by qPCR

RNA was extracted from 100mg liver tissue using 1 ml Trizol reagent. The tissue homogenization was performed using two Tungsten Carbide Beads (Qiagen) (3 mm) on a Tissuelyser II (Qiagen) at a frequency of 30.0 I/sec for 2.5 min. 200ul of chloroform (Sigma-Aldrich) was added and the mixture centrifuged at 12,000 g for 20 minutes at 4°C to separate the aqueous phase which was retained. RNA precipitation was performed using 700µl of isopropanol (Sigma-Aldrich) and centrifuged at 12,000g for 20 min at 4°C and washed 3 times with 80% ethanol. The concentration and purity of total RNA was estimated using a NanoDrop 1000 Spectrophotometer (Thermo Scientific). A QuantiTect Reverse Transcription kit (Qiagen) was used to synthesize first-strand cDNA from 1µg total RNA per sample, with a genomic DNA removal step included, in a total volume of 20ul. The resulting cDNA was diluted 20-fold (working stock) with RNase/DNase free water. qPCR was used to confirm an inflammatory response in the infected fish, using primers targeting the APP encoding genes *saa* and *hamp* (11). For normalization of gene expression, primers targeting *rps13* and *rps29* were used (19). qPCR was performed with 2x SYBR Green I (Invitrogen) master mix on a Mx3005P System (Agilent Technologies). Each assay was run in triplicate using 15µl of reaction mix containing 7.5ul Brilliant III Ultra-Fast SYBR Green (Agilent Technologies), 500nM forward and reverse primers and 5µl of cDNA (2.5ng of total reverse-transcribed RNA). Assays were run with 1 cycle of 95°C for 3 min, followed by 40 cycles of 95°C for 20s and 64°C for 20s. A melting curve (thermal gradient 55°C-95°C) was used to confirm single qPCR products. Each qPCR plate included two no-template (i.e. water) controls. LinRegPCR (20) was used to establish the efficiency of each assay. Gene expression data were analyzed using GeneX 5.4.3 (MultiD Analysis), correcting for differences in efficiency across genes, before target gene expression was normalized to *rps13* and *rps29* and placed on a scale of relative expression.

### snRNA-seq library construction

A protocol adapted from ref (21) was used for nuclear extraction, employing a tween with salt and tris (TST) buffer. Approximately 45mg of each frozen liver sample was placed in a 6-well tissue culture plate (Stem Cell Technologies) with 1 ml TST (2 mL of 2X ST buffer + 120 µL of 1% Tween-20 + 20 µL of 2% BSA brought up to 4 ml with nuclease-free water). The tissue was minced using Noyes Spring Scissors for 10 min on ice. The resulting homogenate was filtered through a 40 µm Falcon cell strainer, and a further 1 mL of TST was added to wash the well and filter. The volume was brought up to 5 mL using 3 mL of 1X ST buffer (diluted from 2xST buffer [292 µl of 146 mM NaCl, 100 µl of 10 mM Tris-HCl pH 7.5, 10 µl of 1 mM CaCl_2_, 210 µl of 21 mM MgCl_2_, brought up to 10ml with nuclease-free water]). The sample was centrifuged at 4°C for 5 min at 500g before the resulting pellet was re-suspended in 1 ml 1X ST buffer and the recovered nuclei were filtered through a 40 µm Falcon cell strainer, Hoechst stained, visually inspected under a fluorescent microscope, and counted using a Bio-Rad TC20. Liver nuclei were processed through the 10X Chromium™ Single Cell Platform using the Chromium™ Single Cell 30 Library and Gel Bead Kit v3.1 and Chromium™ Single Cell A Chip Kit (both 10X Genomics) as per the manufacturer’s protocol. For each sample, the nuclei were loaded into a channel of a Chromium 3’ Chip and partitioned into droplets using the Chromium controller before the captured RNA for each nucleus was barcoded and reverse transcribed. The resulting cDNA was PCR amplified for 14 cycles, fragmented, and size selected before Illumina sequencing adaptor and sample indexes were attached. Libraries were sequenced on a NovaSeq 6000 by Novogene UK Ltd (2×150bp paired end reads).

### Generation of snRNA-Seq count matrix

Raw sequencing data were aligned to the unmasked ICSASG_v2 reference assembly (Ensembl release 104) of the Atlantic salmon genome (22). The analysis was restricted to protein coding genes. Mapping of reads to the genome, assignment of reads to cellular barcodes, and collapsing of unique molecular identifiers (UMIs) was performed with StarSolo v2.7.7a (23). The genome index was generated with standard settings and “*sjdbOverhang*” set to 149. The reads were then mapped with the “STAR” command and following settings: “*soloType=CB_UMI_Simple*”, “*outFilterMultimapNmax = 20*”, “*outMultimapperOrder=random*”, “*soloUMIdedup=1MM_Directional*”, “*soloFeatures GeneFull*”, “*soloBarcodeReadLength = 0*”, “*outFilterMatchNminOverLread = 0*”, “*soloCellFilter = TopCells 100000*”. The top 100,000 cell barcodes ranked by UMI number were retained to ensure the capture of transcriptionally quiet nuclei, lost when using the automated StarSolo filtering algorithm. Mapping statistics for each snRNA-Seq sample is provided in Supplementary Table 1.

### Nuclei filtering and quality control

Nuclei filtering was performed manually on each of the four samples using Seurat v3.1 (24). Ranked barcode plots of UMIs and gene counts were used to identify the lower ‘elbow point’ and cell barcodes with UMI count or gene counts below the elbow point were removed as empty droplets. Further steps were used to identify additional empty droplets: the “*SCTransform”* function (25) was used to normalise the data, prior to centering, scaling, principal component analysis (PCA) and high-resolution clustering (using 30 PCs and resolution of 2). Wilcoxon rank sum differential gene expression tests were used to identify up-regulated genes in each cluster. Cell clusters that both lacked distinguishing markers and had a low median UMI or gene counts (typically 2 median absolute deviations lower than the median across all nuclei) were removed as likely empty droplets or poor-quality nuclei. This process was repeated iteratively on each sample until all such low-quality populations were removed. Likely doublets were identified and removed later in the analysis, after cell identity was established.

### Assignment of cellular identity

Each cell cluster was assigned to one of the major liver cell lineages using *a priori* marker genes (Supplementary Table 2). Populations identified as erythrocytes and thrombocytes were not included in downstream analyses due to their likely origin from contaminating blood. Populations identified as hepatocytes, cholangiocytes, mesenchymal cells, endothelial cells, and immune cells were merged into five separate Seurat objects (using the Seurat “*merge*” function) for separate analyses of each cell lineage. Batch effects across samples were removed in each merged object with Harmony (26). Gene annotations were taken from the Ensembl annotation for Atlantic salmon, and in cases where the gene was not assigned a name, the name of the nearest annotated ortholog in rainbow trout (*Oncorhynchus mykiss*), zebrafish or mouse (*Mus musculus*) was used, or in some cases informed by BLASTp searches against the NCBI non-redundant database.

For each of the five major liver cell lineages retained, data was log normalised, then scaled and centered, before PCA was performed and used as the input to graph-based clustering, using the established Seurat pipeline. The appropriate number of PCs to use for clustering of each sample to minimise technical noise, was determined through visualisations generated through the “*ElbowPlot*” and “*DimHeatmap*” functions. The resolution parameter in the “*FindClusters*” function was tuned to return biologically meaningful heterogeneity with each cell type. At this stage, a further quality control step was performed to remove doublets from each lineage. Specifically, differential gene expression tests were performed and clusters that exhibited canonical markers (Supplementary Table 2) or lineage distinguishing markers (Supplementary Table 3) from two distinct cell lineages yet lacked any unique distinguishing marker genes of their own, were removed as likely doublets, before the data was re-clustered. This process was repeated until no doublet populations remained. For the immune cells, the Seurat object was further split into T cell, Myeloid, NK-like, Neutrophil and B cell objects (see Results) and the same process was repeated (using markers in Supplementary Table 3) and cell sub-clusters identified, before each object was merged back into a single “immune cell” Seurat object. Finally, all five major lineages were merged into a final global liver cell atlas object, with cell identities retained from the cell lineage specific analyses. We used this strategy as opposed to global clustering, as markedly more biologically meaningful heterogeneity could be established in the cell lineage specific analyses.

## Results

### Single-nuclei RNA-Seq atlas of the Atlantic salmon liver

Hepatocytes, the major epithelial cell type within the liver, have been underrepresented in mammalian scRNA-Seq analyses (2), which may be caused by damage occurring during the dissociation step. As Atlantic salmon are ectotherms, we were also concerned that enzymatic dissociation (typically done at >30°C) would activate cell stress and heat shock responses. We consequently decided to generate a snRNA-Seq atlas of salmon liver, using nuclei isolated from freshly flash frozen samples, an approach expected to provide an accurate representation of cell diversity (e.g. 27-28). The profiled liver nuclei were from control fish (n=2) and animals infected by a pathogenic strain of *Aeromonas salmonicida* (n=2), the bacterial agent of furunculosis, a long-standing problem disease in salmonid aquaculture (29). The infected group were sampled 24 hours post *Aeromonas* challenge, previously shown to capture the inflammatory and acute phase response (11). The fish used for sequencing were further selected based on gene expression data using marker genes for the acute phase response (11), which were robustly up-regulated in the infected group (Supplementary Fig. 1).

Across all samples, we generated 47,432 nuclei transcriptomes with median UMI and gene counts per nucleus of 2,105 and 1,065, respectively. This was split across control fish as 11,679 and 11,433 nuclei, and *Aeromonas*-challenged fish as 19,148 and 5,172 nuclei, respectively (Supplementary Table 1). The major cell lineages were identified using a guided graph-based clustering strategy (24), with cell identity assigned using *a priori* defined marker genes (Fig. 1a, b; markers in Supplementary Table 2). Hepatocytes comprised most of the profiled nuclei (88.1%), followed by cholangiocytes (4.3%), immune cells (3.5%), mesenchymal cells (2.6%) and endothelial cells (1.5%) (Supplementary Fig. 2; Supplementary Table 1). Clustering and identification of these major cell lineages was repeatable across individual samples, with each sample contributing a large proportion of the nuclei (Supplementary Fig. 2). Differential gene expression analysis revealed markers for each major liver cell type (Fig. 1c, d; Supplementary Table 3). Marker genes for candidate cell types and sub-populations are hereafter reported according primarily to annotations provided by Ensembl, or in cases where no Ensembl annotation was available, using supplementary BLAST homology searches against the NCBI database to support our inferences. While Ensembl annotation utilizes phylogenetic information to inform homology relationships, it may nonetheless fail to correctly capture orthology of Atlantic salmon genes to mammalian species, particularly for fast evolving and complex gene families. Adding to this challenge, genetic orthology is not a prerequisite for conservation of gene function or expression, which limits our ability to transfer knowledge about cell marker genes from mammalian studies to Atlantic salmon. While the reader must be aware of these caveats, they represent a general issue in functional genomics studies using non-model taxa like salmonids.

**Figure 1.**
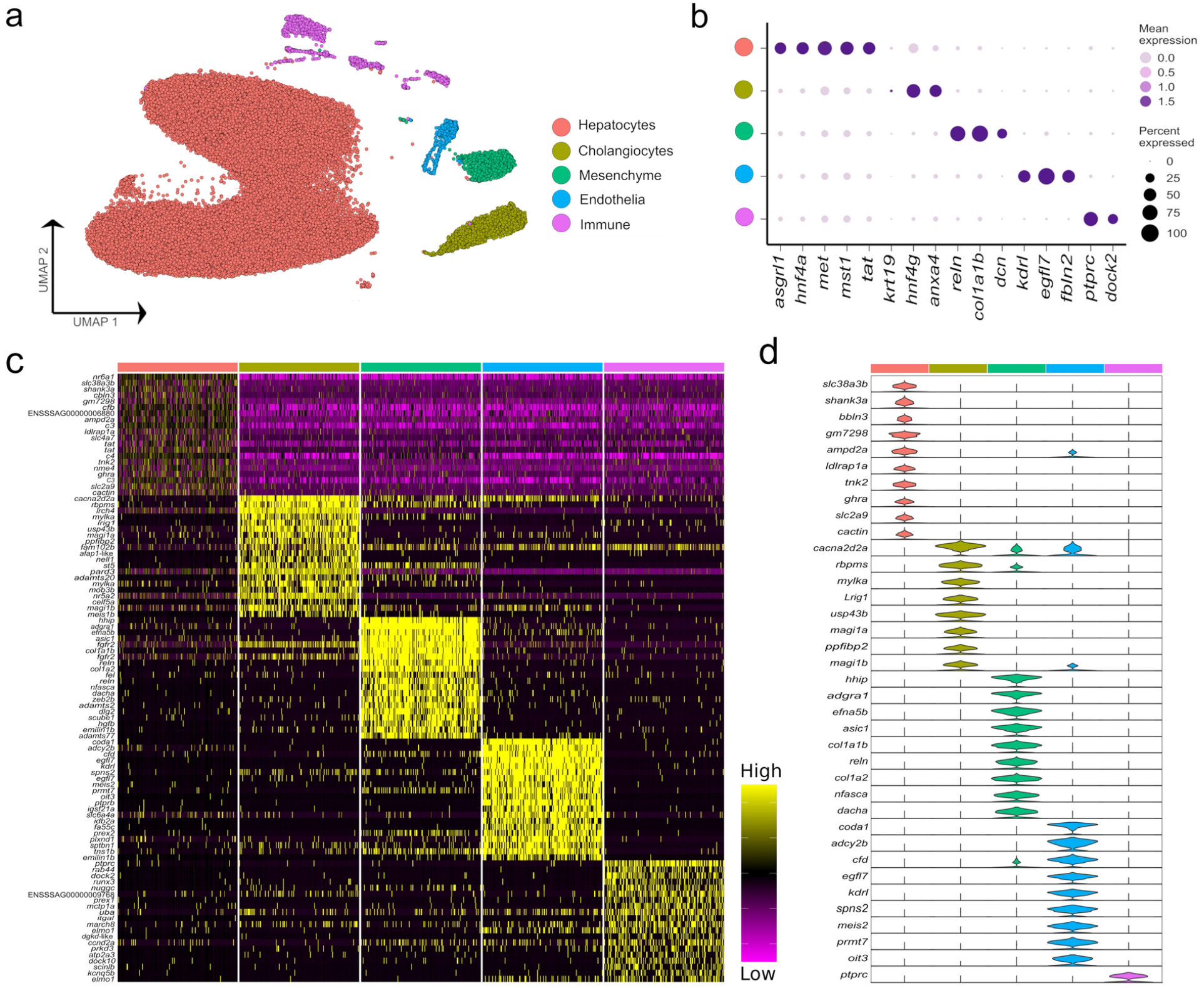
Major cell types in the Atlantic salmon liver defined by 47,982 nuclear transcriptomes. (**a**) UMAP highlighting five main liver cell-type clusters according to *a priori* defined marker genes (see Supplementary Table 2). (**b**) bubble plots showing the expression of *a priori* marker genes the five main liver cell types, including mean expression (bubble intensity) and percentage of nuclei expressing gene (bubble size). (**c**) Heatmap of the top 20 differentially expressed genes per each of liver cell types defined against the background of all other cell types. (**d**) Violin plots demonstrating the expression of the most specific marker genes per each of the main liver cell types (colours are matched to the colours of the 5 cell lineages defined in part (**a**).

While it was possible to identify the five major liver cell lineages of Atlantic salmon using orthologues to marker genes defined in mammalian liver scRNA-Seq datasets (e.g. 6), many markers were notably absent or expressed at very low levels in our snRNA-seq dataset (Supplementary Fig. 3). For example, Atlantic salmon orthologues of the widely used epithelial marker *Epcam* did not show expression in the cholangiocyte cluster (Supplementary Fig. 3). Likewise, salmon orthologues of *Pecam1* and *Pdgfrb*, which are excellent markers of mammalian endothelial and mesenchymal cells, were not detected at significant levels in these cell types in our dataset (Supplementary Fig. 3). Such results may be explained by differences in transcriptome composition between snRNA-seq and scRNA-seq, or by divergence in gene expression between fish and mammals. Differences in expression were also observed between predicted Atlantic salmon orthologues of mammalian marker genes for the major hepatic cell types, with, for example, only one of two *Cdh5* co-orthologues marking the endothelial population, and only one of two *Hnf4a* co-orthologues marking hepatocytes (Supplementary Fig. 3).

Higher resolution clustering captured varying degrees of transcriptomic heterogeneity for each of the five major liver cell types (Fig. 2a, b; Supplementary Table 4), which was consistent across the four samples (Supplementary Fig. 4). We observed a split of hepatocytes into sub-populations explained largely by infection status, with 75.9% of ‘control-associated’ hepatocytes deriving from control fish and 73.3% of ‘infection-associated’ hepatocytes deriving from *Aeromonas*-challenged fish (expanded in next section). Significant markers for the latter were dominated by genes encoding APPs, many of which were observed to be expressed across all cell types (Fig. 2b). This is likely a consequence of the numerically dominant hepatocytes (Fig. 1a) leaking mRNA from highly expressed genes into the ambient RNA. Similar examples of this type of leakage can be observed in many liver scRNA-Seq datasets with, for example, hepatocyte expressed genes *Alb* and *Hp* being widely ‘expressed’ in non-hepatocyte cells (e.g. 3, 30).

**Figure 2.**
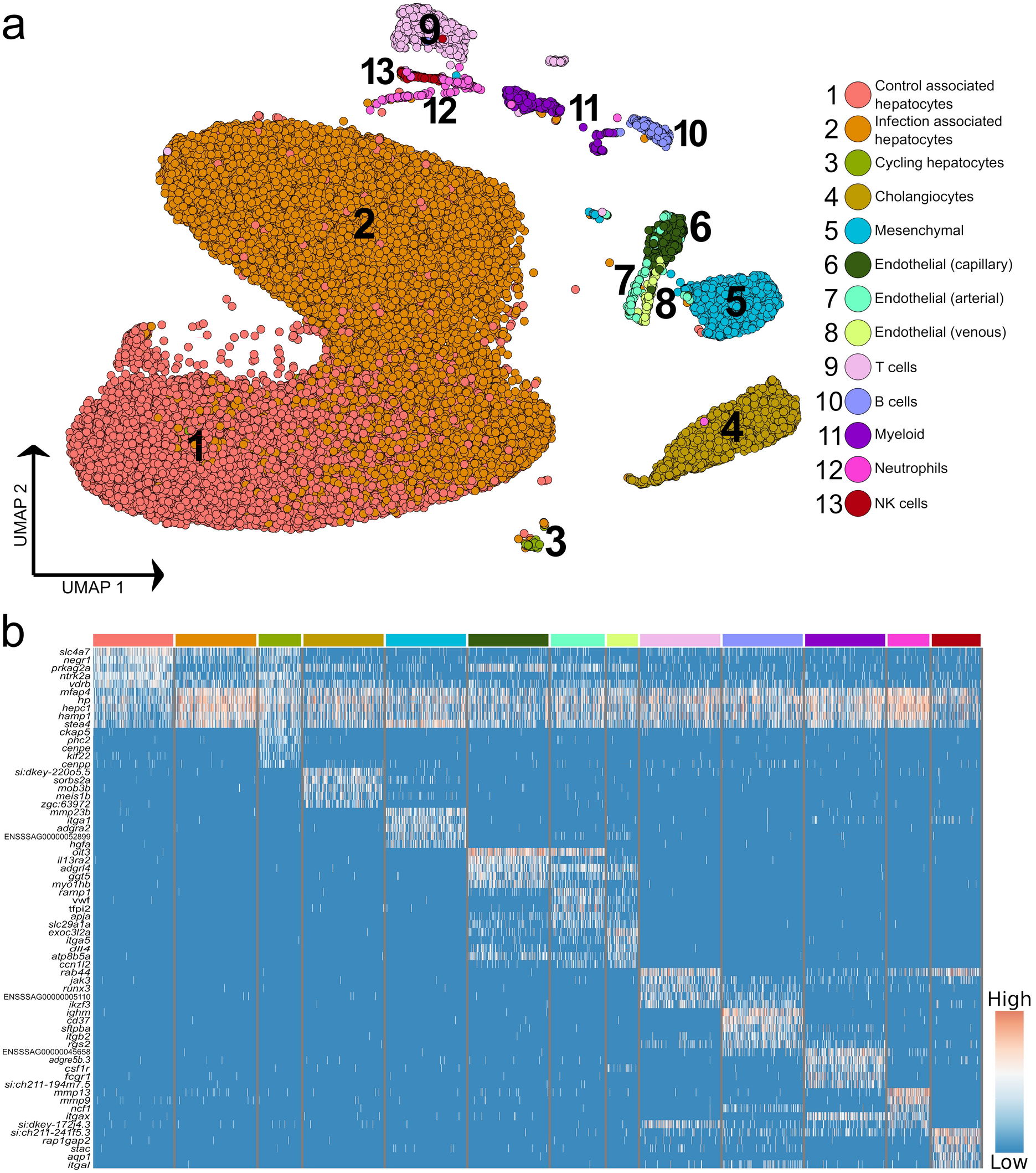
Higher resolution atlas of Atlantic salmon liver cells defined by snRNA-Seq. (**a**) Unbiased graph-based clustering reveals varying heterogeneity across the major liver cell types, presented on a UMAP. Each cell population retains the gene signature of the parent lineage (Supplementary Fig. 2), while also displaying its own distinct transcriptomic profile, presented here as a heatmap (**b**), inclusive of the top 10 marker genes based on differential gene expression against all other cell clusters. The colour bars above columns on the heatmap illustrate the cell types to which the genes shown were identified as markers (matched to part a).

The immune compartment contained identifiable T and B cells, along with candidate populations of neutrophils, myeloid cells, and NK-like cells (Fig. 2b). The transcriptome of cholangiocyte nuclei was homogeneous, while limited heterogeneity was identified in the endothelial and mesenchymal cells (Fig. 1a; Supplementary Fig. 5 and 6). The endothelia sub-clusters had a clear biological interpretation, with Atlantic salmon orthologues to marker genes from mammals distinguishing arterial and venous derived endothelial cells (31) (Supplementary Fig. 5, marker genes in Supplementary Table 5). However, the mesenchymal sub-clusters were not readily biologically interpretable (Supplementary Fig. 6; Supplementary Table 6).

### Hepatocyte remodelling dominates the liver response to bacterial infection

To explore how hepatocyte heterogeneity contributes to the response to *Aeromonas* infection, we analysed 41,792 available hepatocyte nuclei transcriptomes. Clustering using the most variable genes in this compartment identified nine sub-populations (H1-H9) (Fig. 3a; marker genes in Supplementary Table 7), with several showing marked differences in abundance between control and infected fish (Fig. 3b). H1-H4 comprised 90.2% of hepatocyte nuclei, with H1 and H2 deriving mainly from control fish and showing highly correlated transcriptomes (Fig. 3b, c). H3 and H4 comprised 70.0% of nuclei from infected fish and showed closely related transcriptome profiles (Fig. 3b, c).

**Figure 3.**
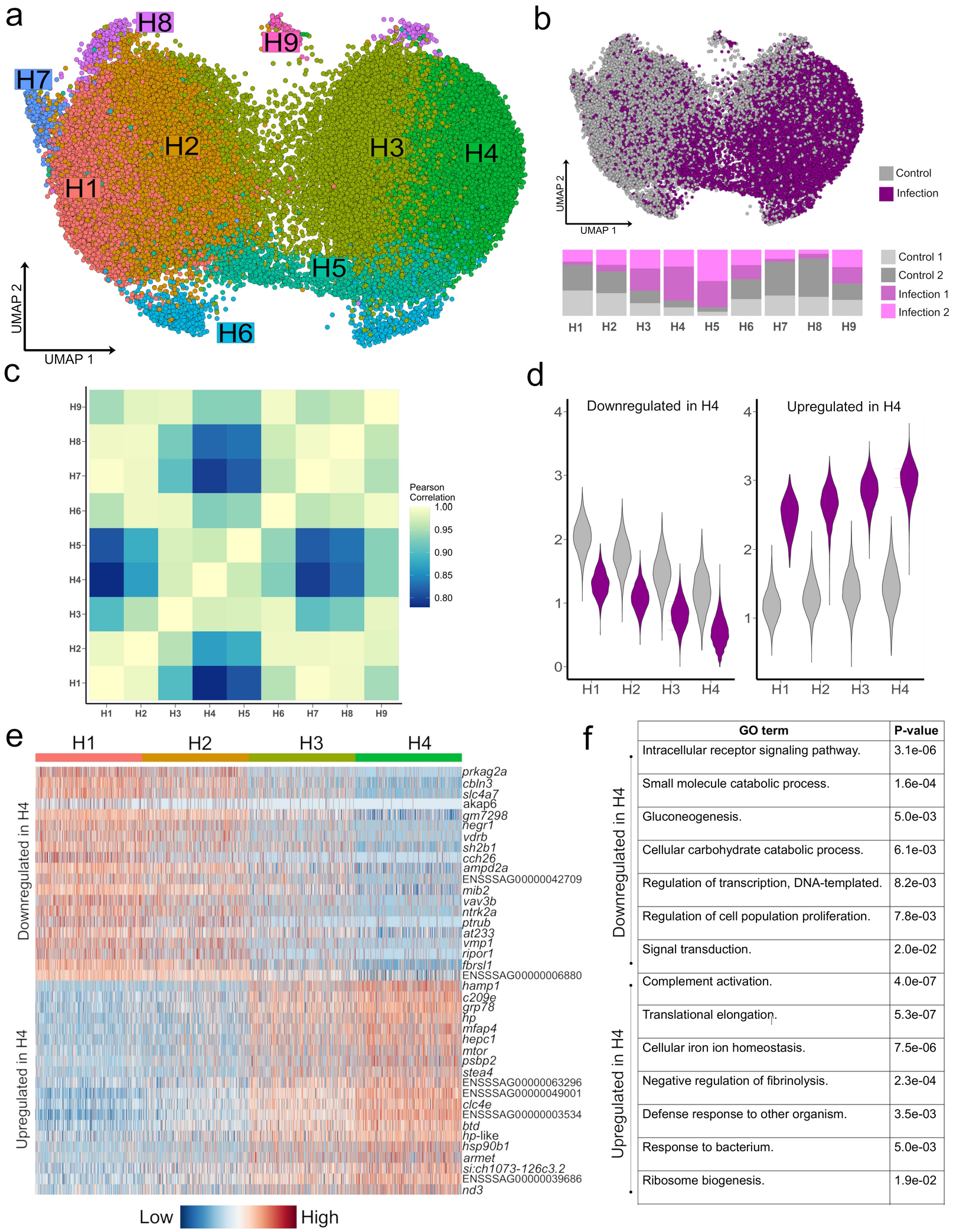
Striking remodeling of the hepatocyte transcriptome in response to bacterial infection. (**a**) UMAP visualisation of 41,792 hepatocyte nuclei, with sub-clustering performed using the most variable genes restricted to this cell lineage. (**b**) Shows the same UMAP with nuclei coloured by infection state (top) and the proportion of nuclei originating from each sample after normalising for different nuclei numbers across samples (bottom). (**c)** Pearson correlation of the expression values for the top 2,000 most variable genes across the nine hepatocytes populations. (**d**) Violin plots of mean expression for the 20 most down-regulated (left) and up-regulated (right) genes based on log-fold change in H4 vs. H1. (**e)** Heatmap of the top 20 most upregulated and top 20 most downregulated genes in H4 relative to H1, illustrating a gradient of expression from H1 → H2 → H3 → H4 (**f**) Example enriched GO terms in H4 based on all up-regulated and all down-regulated genes in H4 vs. H1 (full data provided in Supplementary Tables 10 and 11)

Hepatocyte nuclei derived from infected fish increased from H1 (16.9%), to H2 (28.2%), to H3 (64.3%) to H4 (80.8%), with 2,842 genes differentially expressed on this gradient (Fig. 3d, e; Supplementary Tables 8 and 9). 379 genes were up-regulated in H4 vs. H1, showing overrepresented functions linked to host defense and the acute phase response (‘*complement activation’*, ‘*defense response to other organism’*, and ‘*cellular iron homeostasis*’), in addition to translational processes (e.g. ‘*translational elongation’*) (Fig. 3f, Supplementary Table 10). This response was dominated by genes encoding APPs including hepcidin, haptoglobins, ferritins, transferrin, fibrinogens, ceruloplasmin, angiotensinogen, serum albumins, apolipoproteins, and c-reactive protein (Supplementary Table 8). One of the top up-regulated genes (3.7-fold up-regulated; ENSSSAG00000046715) encodes mechanistic target of rapamycin kinase (mTOR), a master regulator of translation (32). mTOR biasedly promotes translation of ribosomal protein genes (32), which is notable as 28 such genes, encoding many proteins comprising the large and small ribosomal subunits of salmonid fish (33), were up-regulated in H4 vs. H1, with only one downregulated (Supplementary Tables 8 and 9). We further observed up-regulation of *copb1* (ENSSSAG00000068484), *grp78* (ENSSSAG00000054661), *sec61a1* (ENSSSAG00000039008), and *srprb* (ENSSSAG00000046456), encoding major components of the COPI and translocon complexes, representing key protein secretion pathways (34).

H5 nuclei were mainly from infected fish (Fig. 3b) and showed highly correlated transcriptomes to H3/H4 (Fig. 3c), sharing many of the same key markers up-regulated in H4 vs. H1, but also specific markers (Supplementary Table 7) associated with NF-κβ signaling, including *relb* (ENSSSAG00000052551), encoding a component of the NF-κβ transcription factor complex. This is notable, as H5 also showed the highest expression among all hepatocyte sub-populations of *stat3* (ENSSSAG00000003657), encoding signal transducer and activator of transcription 3, which acts downstream of NF-κβ in mammalian hepatocytes to activate inflammation driving the acute phase response (35).

2,278 genes were downregulated in H4 vs. H1 (Supplementary Table 9), enriched in functions related to signaling (e.g. *‘intracellular receptor signaling pathway*’), metabolism (e.g. ‘*gluconeogenesis*’), cell differentiation (e.g. ‘*stem cell differentiation*’) and transcription (e.g. ‘*regulation of transcription, DNA-templated’*) (Fig. 3f) (Supplementary Table 11). The top downregulated gene was *prkag2a* (4.0-fold down-regulated; ENSSSAG00000079550), encoding a subunit of the AMPK complex - a master regulator of metabolism including gluconeogenesis (36). Also downregulated were master hepatic transcription factors for lipid and glucose metabolism pathways that interact with AMPK (36), including genes encoding hepatocyte nuclear factor 1 (HNF1) (ENSSSAG00000006158) and HNF4 (ENSSSAG00000047055) (37), carbohydrate-responsive element-binding protein (ENSSSAG00000039257) (38), and forkhead box proteins O1 / O3 (ENSSSAG00000054669 / ENSSSAG00000055241) (39) (Supplementary Table 9). Further evidence for repression of anabolism included downregulation of genes encoding growth hormone receptor (GHR) (ENSSSAG00000065355 and ENSSSAG00000081526) and Stat5a/b (ENSSSAG00000010616, ENSSSAG00000003584 and ENSSSAG00000048873), which act downstream of GHR to activate growth and cell proliferation genes (40). In addition, a gene was downregulated encoding glucocorticoid receptor (GR) (ENSSSAG00000062169), which promotes expression of gluconeogenesis genes, while its interaction with Stat5 is required for transcriptional activation of growth genes via GHR signalling (40).

### Additional hepatocyte heterogeneity includes polyploid cells

The remaining hepatocyte subclusters were not linked to infection status (Fig. 3b). H7 likely represents hepatocytes that have undergone polyploidization, which occurs progressively during aging in mammals, such that 4n-16n cells make up a large fraction of liver cells by adulthood (41). Polyploidy in H7 is indicated by a striking concordance of genes representing highly specific markers for H7 and those shown elsewhere to be up-regulated in 4n vs. 2n mammalian cells, encoding DNA primase subunit 2 (ENSSSAG00000001875), replication protein A 70 kDa DNA-binding subunit (ENSSSAG00000050927, ENSSSAG00000050209) and DNA polymerase (ENSSSAG00000078390), among others (42). The close relationship of H7 to H1 (Fig. 3a, c) implies that polyploid hepatocytes derive from those supporting routine metabolism.

H9 showed many specific markers encoding mitosis proteins, e.g. cytoskeleton-associated protein 5 (ENSSSAG00000066206), centromere protein E (ENSSSAG00000073454), abnormal spindle-like microcephaly-associated protein homolog (ENSSSAG00000053226) and kinesin family member 23 (ENSSSAG00000044381), indicating these are cycling hepatocytes. H6 expressed a small number of highly specific markers, including genes encoding ligand of numb-protein X1 (*lnx1*) (ENSSSAG00000072728 and ENSSSAG00000070275), a E3 ubiquitin ligase that targets a wide range of proteins, including CD8 expressed on T-cells in mammals (43). H8 was biasedly represented by control fish and most correlated with H1/H2 in transcriptome profile (Fig. 3c), expressing marker genes described as ‘novel’ in the Ensembl annotation (Supplementary Table 7).

### Immune cell heterogeneity in Atlantic salmon liver

3.4% (n=1,620) of the liver nuclei were derived from immune cells (Fig. 4a, b). Combinations of canonical marker expression was used to classify T cells (*cd3e, tox2* and *tcf7*), B cells (*ighm, cd37, cd79a*), NK-like cells (*prf1*.*3* and *runx3* with absence of *Cd3*e) and myeloid cells (*mpeg1, cd63, csf1r, lyz2*) (Fig. 4b; marker genes provided in Supplementary Table 12). We also identified a candidate population of neutrophils based on marker genes that showed highest specificity or expression for neutrophils among different immune cells in the human protein atlas, namely *itgax* (ENSSSAG00000049715), *ncf1* (ENSSSAG00000079828), and *mmp9* (ENSSSAG00000069874) (Fig. 4b). *Mmp9* was an effective marker for neutrophils in other fish and has a role in driving neutrophil migration in mammals (44,45). The NK-like cells represent a tentative annotation owing to a lack of certain markers for NK cells, i.e. genes encoding granzymes. Each immune cluster had substantial contributions from all samples, except the candidate neutrophils, which derived mainly from one *Aeromonas*-challenged fish (Supplementary Fig. 4). Sub-clustering revealed heterogeneity in the T and myeloid cells, but not the NK-like cells, B cells (with no evidence of plasma cells) or candidate neutrophils.

**Figure 4.**
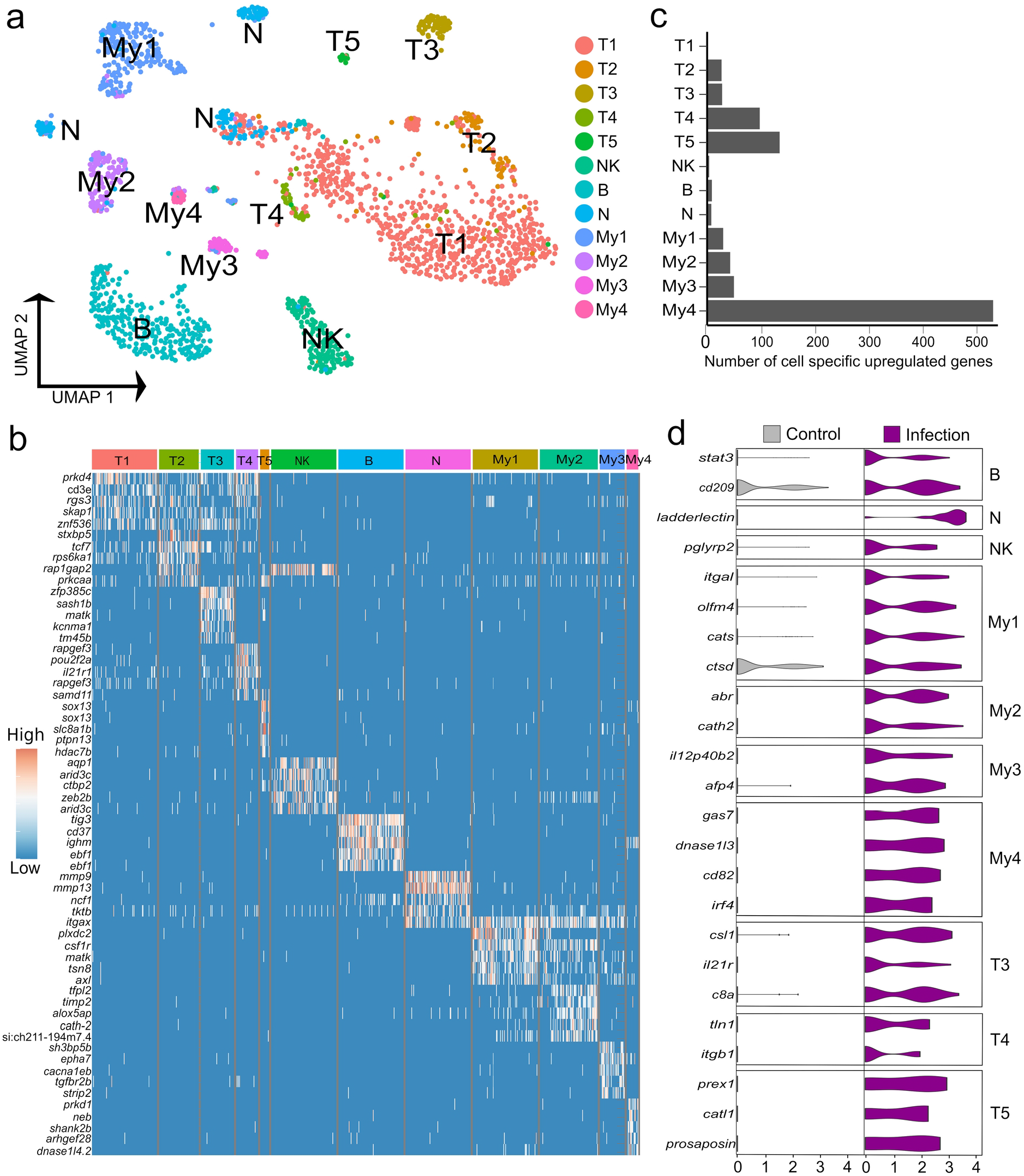
Heterogeneity in Atlantic salmon immune cells. (**a**) UMAP visualization of 1,620 immune nuclei. (**b**) Heatmap of top 5 markers genes for each immune sub-cluster relative to all other liver immune cells, sorted by log2 fold change. (**c**) Number of cell-specific genes up-regulated by infection in immune sub-clusters (**d**) Examples of genes showing cell-specific up-regulation in response to *Aeromonas* infection across the breath of immune cell heterogeneity identified.

Five T cell sub-clusters were compared using differential expression tests (Fig. 4a; Supplementary Table 12). Only T1 and T2 expressed *cd4* genes to low levels, specifically ENSSSAG00000076631, encoding CD4-1, which is also expressed by macrophages, and ENSSSAG00000076595, encoding CD4-2; shown elsewhere to be expressed by all CD4^+^ T cells (46). We did not identify any *CD8* expressing T cells, likely due to low expression levels. T1 was the largest sub-cluster but showed few specific markers. T2 expressed genes encoding receptors involved in T cell activation. This included *cd28* (ENSSSAG00000060163), encoding the main co-stimulatory T cell receptor (47) and *cd44* (ENSSSAG00000076128), a widely used T cell adhesion, co-stimulation and activation marker (48). However, T2 cells did not specifically express any genes annotated as *ctla-4*, an IgSF member induced during T cell activation that regulates CD28 activity. Furthermore, T1 and T2 both expressed *tcf7* (ENSSSAG00000006857) at a much higher level than T3-T5, encoding the master Wnt pathway transcription factor, which is most highly expressed on naïve mammalian T cells (49) and was a specific marker for resting CD4^+^ T cells in humans (50). T1 and T2 also expressed *foxp1b* (ENSSSAG00000077820) at a higher level than T3-5; a gene essential for quiescent naive T cells in mammals (51). T1 and T2 therefore appear to be constituted mainly of resting T cells. T3 appears to be an activated T cell population based on specific expression of *slamf1* (ENSSSAG00000043093) encoding *CD150* (52). T3 expressed many highly specific markers, including the integrin coding gene *itgal* (ENSSSAG00000046537; encoding CD11a) and *itgb2* (ENSSSAG00000022772; encoding CD18), whose products form lymphocyte function-associated antigen 1 (LFA-1), a molecule with key roles in T cell activation and migration, in addition to cytotoxic and memory responses (53). The human orthologue of a highly expressed T3 specific marker, *cacna2d2a* (ENSSSAG00000071299), encoding a calcium voltage-gated channel, was not detected in any immune cell in the human protein atlas, implying a teleost-specific T cell marker. T4 expressed several activation markers including *pou2f2a* (aka *oct2*) (ENSSSAG00000071136) (54), *CD226* (55), along with two distinct *ctram* genes (encoding cytotoxic and regulatory T-cell molecule), previously shown in mammals to be required for differentiation of cytotoxic CD4^+^ T cells (56). T5 specifically expressed two paralogues of *sox13* (ENSSSAG00000077869, ENSSSAG00000058488), the defining vertebrate transcription factor for the γδ T lineage (57).

My1 and My2 markers are associated with monocytes and macrophages (Supplementary Table 12). My1 specifically expressed *cd4-1* (ENSSSAG00000076631), at a level higher than any T sub-cluster, likely representing a phagocytic CD4^+^ macrophage characterized in rainbow trout (46). My2 expressed specific monocyte marker genes including *csf3r* (ENSSSAG00000041566) and *timp2* (ENSSSAG00000042353 and ENSSSAG00000064056). Two *csf1r* copies were identified with reciprocal higher expression in My1 (ENSSSAG00000004088) and My2 (ENSSSAG00000061479). High *flt3* (ENSSSAG00000009390) expression in My4 supports an annotation as dendritic cells (DCs) (58), with specific up-regulation of *cd9* (ENSSSAG00000059637) and *lamp2* (encoding CD107b) (ENSSSAG00000074801) genes, consistent with monocyte-derived DCs in mammals (59-60). My3 expressed the second highest level of *flt3*, while the top My3 marker gene, *ptprsa* (ENSSSAG00000051752), is a specific marker for plasmacytoid DCs (pDCs) in mice and human (61). The relationship of My3 to pDCs is also supported by specific expression of *sbf1* (ENSSSAG00000071635) (62). However, another My4-specific marker gene, *tcf4* (ENSSSAG00000071044), encodes a transcription factor required for pDC development (63).

### Immune cell-specific responses to Aeromonas insult

To uncover the role of hepatic immune cell heterogeneity in the response to *Aeromonas* challenge, we performed differential expression tests comparing nuclei from control and infected fish for each immune sub-cluster (Fig. 4c, d). 819 genes showed significant up-regulation (criteria: P < 0.05; Log2FC > 1), among which 274 (33%) and 72 (8.8%) were up-regulated by most (≥7 of the 12 sub-clusters) or all immune cell types, respectively, and 271 (33%) showed immune sub-cluster specific up-regulation (Fig. 4c; Supplementary Table 13).

B cells specifically up-regulated two genes, namely *stat3* (2.4-fold up-regulated; ENSSSAG00000003657), which is essential for B cell differentiation (64) and a gene annotated zgc:174904 (2.1-fold up-regulated; ENSSSAG00000070511), encoding a 304 amino acid protein with a CD209/DC-Sign-like, C-type lectin-like domain (InterPro domain: IPR033989). C-type lectin/DC-SIGN is a broad-specificity PRR that detects bacteria by binding mannose or carbohydrate structures (65). In the candidate neutrophils, the top up-regulated gene was *ladderlectin* (38.9-fold up-regulated; ENSSSAG00000039613), encoding a soluble lectin that bound *Aeromonas* in salmonids, leading to bacterial killing actions (66). The NK-like cells up-regulated few genes specifically, one of which was *pglyrp2* (2.3-fold up-regulated; ENSSSAG00000054105), a peptidoglycan recognition protein with enzymatic activity targeting and limiting the inflammatory effects of bacterial peptidoglycan (67).

Among the T cell populations, T3, T4, and the γδ T cells (T5), showed the strongest responses to *Aeromonas* infection. The top up-regulated genes in T3 included *csl1* (4.3-fold up-regulated; ENSSSAG00000004327), encoding a L-rhamnose-binding lectin that binds bacteria and enhances phagocytosis in salmonids (68), and *c8b* (4.1-fold up-regulated; ENSSSAG00000073702), encoding a core component of the complement membrane attack complex. In T4, up-regulated genes included *tln1*, encoding Talin-1 (4.8-fold up-regulated; ENSSSAG00000063331), which is known to regulate the integrin LFA-1 complex (defining T3; see last section), and is required for sustained interactions between APCs and T cells, as well as T cell proliferation (69). T4 also up-regulated *itgb1* (encoding CD29, also called β1-integrin) (4.8-fold up-regulated; ENSSSAG00000007621), a signature marker for cytotoxic T cells in humans (70). In T5, among the top-up-regulated genes was *prex1* (4.8-fold up-regulated; ENSSSAG00000044871), a signaling molecule that promotes expression of key interleukin cytokines in activated human T cells, including IL-2 (71). T5 also up-regulated *catl1*, encoding cathepsin L (8.0-fold up-regulated; ENSSSAG00000077309), which regulates T cell cytotoxicity (72) and an unannotated gene encoding a protein with saposin-like domains, which is annotated as *prosaposin-*like in NCBI (7.4-fold up-regulated; ENSSSAG00000009411). In mammals, *prosaposin* encodes the precursor to all saposin lysosomal proteins, which are known to have antibacterial activity and play a key role in presenting lipid antigens to CD1-restricted T cells (73).

My1 specifically up-regulated 17 genes, including a different *itgal* paralogue (encoding CD11A) to that noted as a marker for T3 (4.2-fold up-regulated; ENSSSAG00000046996), encoding a component of LFA-1 essential to the immune response of mice to *Mycobacterium tuberculosis*, supporting T cell-mediated activation and recognition of infected macrophages (74). My1 also specifically up-regulated *olfm4* (4.1-fold upregulated; ENSSSAG00000046003), encoding a glycoprotein induced in mice macrophages by *Helicobacter pylori* infection, which regulates inflammatory responses (75). My1 further specifically up-regulated *cats* (ENSSSAG00000070942) and *ctsd* (ENSSSAG00000027269), encoding cathepsin S and D, proteolytic enzymes with established macrophage roles in bacterial killing and antigen processing (76). My2 specifically up-regulated 32 genes, including *cath2* (6.6-fold upregulated ENSSSAG00000049319), encoding an antimicrobial peptide that increased in abundance in response to *Aeromonas* infection in salmon plasma (77) and *abr* (6.6-fold upregulated; ENSSSAG00000080204), encoding a GTPase-activating protein that down-regulates the inflammatory actions of macrophages (78).

My3 specifically up-regulated 16 genes, including *IL12p40b2* (5.7-fold up-regulated; ENSSSAG00000069633), best known as a component of IL-12 and/or IL-23 heterodimers, but that also has defined cytokine functions as a monomer protein, including promotion of DC migration in response to bacterial infection in mammals (79,80). Another induced gene is *afp4* (3.7-fold up-regulated; ENSSSAG00000072959), encoding type IV ice-structuring protein LS-12, an apolipoprotein-like molecule that increases in abundance dramatically in salmon plasma following *Aeromonas* infection (81). My4 specifically up-regulated 64 genes, including *Gas7* (8.56-fold up-regulated; ENSSSAG00000076110), which has a crucial role in phagocytosis (82), *dnase1l3* (8.4-fold up-regulated; ENSSSAG00000066441), which controls inflammasome-induced cytokine secretion (83), and *CD82* (6.9-fold up-regulated; ENSSSAG00000052206), previously shown to be up-regulated during DC activation, where is promotes stable interactions between DCs and T cells, and MHC-II maturation (84). My4 further upregulated *irf4* (6.28-fold up-regulated; ENSSSAG00000039730), a gene essential to the ability of DCs to promote Th2 differentiation and inflammation (85).

## Discussion

This study greatly enhances our knowledge of liver function in a salmonid fish with global commercial and scientific importance. The major advancement compared to previous work comes from our application of snRNA-Seq, which, in contrast to previously past bulk transcriptomic or proteomic studies, allowed us to identify heterogeneity within multiple hepatic cell populations, before dissecting its role in host defense following bacterial infection. Furthermore, a plethora of novel marker genes are reported for developmentally and functionally diverse hepatic cell types, which will be useful for future studies investigating traits relevant to salmonid health and immunological status.

Our results demonstrate the essential contribution of hepatocytes to the antibacterial and acute phase response in Atlantic salmon. Transcriptomic heterogeneity in the dominant hepatocyte population (i.e., H1-4; Fig. 3) was inconsistent with distinct hepatocyte populations. Instead, our data supports a single hepatocyte population that can exist in radically distinct transcriptional states dependent on infection status. At one extreme are the hepatocytes that dominate the liver of healthy fish (H1/2), which appear to be performing routine functions controlled by master hepatic transcription factors and signaling pathways. Conversely, the liver of bacterially infected Atlantic salmon was dominated by hepatocytes (H3/4) that downregulated master hepatic pathways (e.g. controlling metabolic functions), and up-regulated a suite of genes encoding APPs and innate immune molecules. This includes many APPs routinely detected in Atlantic salmon plasma by proteomics (86) implying extremely high abundance. As most plasma proteins derive from liver, this aligns with our finding that these ‘defense-specialized’ hepatocytes strongly up-regulated mTOR, its target ribosomal protein-coding genes, and genes from protein secretion pathways, presumably to boost APP translation and secretion rates during the acute phase response. This striking repurposing of hepatocyte function upon infection illustrates the vital role these cells play as a hub for cross-talk between metabolism and immunity, presumably allowing energetic resources to be allocated towards clearing a pathogen at the short-term cost of limiting investment into routine functions e.g. supporting growth (14).

Interestingly, a subpopulation of defense-specialized hepatocytes (i.e. H5; Fig. 3), almost exclusively derived from infected fish, specifically expressed genes associated with NF-κβ signaling (e.g. *relb* was ∼10-fold up-regulated compared to the average across the eight other hepatocyte subpopulations) and also up-regulated *Stat3*, which is indispensable for activation of APP and protein secretion pathway gene expression during bacterial infection in mice, acting downstream of NF-κβ (35, 87). As APP and secretory protein pathway genes were strongly up-regulated in the dominant subpopulations of defense-specialized hepatocytes (H3/4), which lacked significant *relb* expression, H5 may represent an intermediate hepatocyte state, where the activation of the APP response and associated secretory pathway is first initiated. A previous scRNA-Seq study of zebrafish liver failed to identify hepatocytes showing any equivalent specialization towards host defense (18), while another identified a minor hepatocyte population enriched for immune functions (17), which may be analogous to H3-H5. Differences with these past zebrafish studies may reflect the fact that both studies utilized control zebrafish lacking immune stimulation. However, the fact that H3-5 comprised a significant fraction of hepatocytes in our control fish could also be explained by differences in liver function, potentially indicative of a higher baseline inflammatory state in the liver of the Atlantic salmon population we studied.

While representing a small population of liver nuclei, we also offer evidence for polyploid hepatocytes in Atlantic salmon. The functional role of polyploidy in mammalian liver remains ill-defined, despite extensive study over decades (41). Work done over 40 years ago showed that the liver of several teleosts contained polyploid hepatocytes (88), so our result is perhaps not unexpected. Polyploidy increases with aging in mammals (41), which may explain why this hepatocyte population was so limited in the fish used in our study, which were juveniles. However, polyploid hepatocytes were not reported in past scRNA-Seq studies of zebrafish liver, which included an 18-month-old adult population (17), representing half the adult lifespan for this species. More work is required to understand the role of hepatocyte polyploidy in teleost health and disease.

It is important to acknowledge that comparing our results with liver scRNA-Seq studies in other species has challenges due to fundamental differences in experimental execution, which, for example, is linked to striking differences in the composition of the cells captured. For example, a preprint by ref (18) surprisingly identified T cells as the dominant liver population, with hepatocytes reflecting a smaller proportion than expected. While underrepresentation of hepatocytes has been observed in several mammalian liver scRNA-Seq studies, a separate zebrafish scRNA-Seq study identified hepatocytes as the most abundant liver cell type (17), albeit at a smaller fraction than for our snRNA-Seq atlas in Atlantic salmon. This perhaps illustrates a recognized benefit of snRNA-Seq compared to scRNA-Seq; a more accurate representation of the true tissue cell diversity (e.g. 27-28, 89). Considering our limited knowledge of cellular diversity in most fish species, including salmonids, careful comparisons of results from scRNA-Seq and snRNA-Seq will be required to establish baseline expectations for future studies.

This is the first single cell transcriptomic study to characterize immune cell heterogeneity in the liver of a salmonid fish, and the first single cell study in any teleost to characterize transcriptomic responses of specific hepatic immune cell subtypes to infection. Past scRNA-Seq studies of zebrafish liver painted very distinct pictures of lymphocyte diversity, with one reporting no B cells, and little T cell heterogeneity (17). Conversely, a recent preprint reported a small B cell population, and six T cell sub-clusters capturing distinct CD8 and CD4 subsets (18). While we also identified a single B cell population and multiple T cell sub-clusters, the identity of T cells was markedly less clear in our data, due partly to a general lack of *CD4* and *CD8* expression, which may reflect a limitation of snRNA-Seq, or the lower sequencing depth in our study compared to ref (18). However, unlike these previous studies, we identified a small population of *Sox13*^+^ γδ T cells, an ancient vertebrate T cell subtype with roles bridging adaptive and innate immunity. In zebrafish, γδ T cells possess phagocytic ability and act as APCs that activate CD4^+^ T cells, inducing B cell proliferation (90). In mammals, γδ T cells increase dramatically in liver during inflammatory conditions (91) and produce IL-17 essential for the innate response to bacterial infection (90). While γδ T cells have not been reported among the plethora of immune cells reported to date in scRNA-Seq studies spanning different teleost species (93), we find them readily identifiable in Atlantic salmon by *Sox13* expression in multiple tissues (not shown). Consistent with their known functions, salmon liver γδ T cells up-regulated genes with roles spanning innate and adaptive immunity during the early response to *Aeromonas* infection.

Our study also identified evidence for myeloid heterogeneity within the Atlantic salmon liver, including two candidate DC populations that showed a strong response to bacterial infection, up-regulating genes required for interaction with T cells, phagocytosis and inflammasome activation, suggesting conserved roles between mammals and salmonids, as shown elsewhere (94). DCs were not reported in recent scRNA-Seq studies of zebrafish liver (17-18) but were reported in Atlantic cod spleen (92). We also observed two distinct candidate macrophage populations, matching the level of heterogeneity recently reported in zebrafish (17-18). A natural question relates to the relationship of teleost macrophages and mammalian Kupffer cells. Past histology work largely agrees that Kupffer cells are amniote-specific and hence not present in teleost liver (96), including for Atlantic salmon (97). Consistent with this notion, a recent study in zebrafish identified that resident macrophages were not located in the sinusoidal space, and lacked phagocytic ability (98). Hence, while Kupffer cells were not expected among our identified macrophage populations, it is worth noting the expression of the (Ensembl) predicted salmon orthologues of mammalian *CLEC4F* (ENSSSAG00000040735 and ENSSSAG00000076658). *CLEC4F* encodes a C-Type Lectin (also known as Kupffer cell receptor) expressed specifically in Kupffer cells to the exclusion of non-liver macrophages (99; 100). In our Atlantic salmon dataset, one predicted orthologue of mammalian *CLEC4F* is expressed across all four myeloid populations (ENSSSAG00000076658), showing highest expression in My2, whilst the other (ENSSSAG00000040735) was expressed across all immune cell types (Supplementary Table 11). Furthermore, both genes were among the top up-regulated genes in hepatocytes following *Aeromonas* infection (Supplementary Table 8). These findings highlight the potential for striking differences in marker gene cell-specific expression and presumably function for homologous genes shared by mammals and teleosts.

Having robust marker genes for different cell-types is clearly essential to accurately study the cell biology of any species. However, single cell transcriptomics studies performed to date clearly demonstrate that marker genes can vary markedly across species, especially for immune cells. (93, 101). This demands a broader uptake of single cell transcriptomics in more species to define conserved from non-conserved marker genes, and to separate true biological or evolutionary differences in cell heterogeneity (and associated marker gene expression) from differences introduced by technical reasons discussed earlier. In salmonid fishes, the presence of an ancestral whole genome duplication, and the associated retention of many duplicated genes that have diverged extensively in tissue expression (22, 102-104), further challenges the transfer of knowledge on marker genes from model species - more work is required in this area. In summary, our comprehensive dissection of the Atlantic salmon liver using single cell transcriptomics has generated numerous species-specific marker genes for a range of immune and non-immune cell types that contribute to health and immunological traits of relevance to sustainable aquaculture. Our results can further be used to extract cell-specific information from both existing and future bulk gene expression studies.

## Author contributions

Designed study: DJM, RRD, RST; coordinated sampling and disease challenge experiment: SAM; performed fish sampling: SN, SAM, DJM; performed quantitative PCR: SN; optimized nuclear isolation: RRD; generated snRNA-Seq libraries: RD, RRD; provided infrastructure for snRNA-Seq: NCH; performed bioinformatics: RST; interpreted immunological data: TCC, PB; drafted manuscript: DJM, RST, RRD; made figures and tables: RST, DJM; contributed to data interpretation and finalization of manuscript: all authors.

## Acknowledgements

This study was funded by grants from the Scottish Universities Life Sciences Alliance (Technology Seed Funding Call), the University of Edinburgh’s Data Driven Innovation Initiative (Scottish Funding Council Beacon ‘Building Back Better’ Call), and the Biotechnology and Biological Sciences Research Council, including the institutional strategic programme grants BBS/E/D/10002071 and BBS/E/D/20002174 and the responsive mode grant BB/W005859/1. NCH is supported by a Wellcome Trust Senior Research Fellowship in Clinical Science (ref. 219542/Z/19/Z). For the purpose of open access, the author has applied a Creative Commons Attribution (CC BY) licence to any Author Accepted Manuscript version arising from this submission.

## Data Availability

All data generated in this study has been made available in the GEO database (https://www.ncbi.nlm.nih.gov/geo/). Accession: GSE207655.

