## Supplementary Figures 1-7 for "Single cell transcriptomics of Atlantic salmon (*Salmo salar* L.) liver reveals cellular heterogeneity and immunological responses to challenge by *Aeromonas salmonicida*"

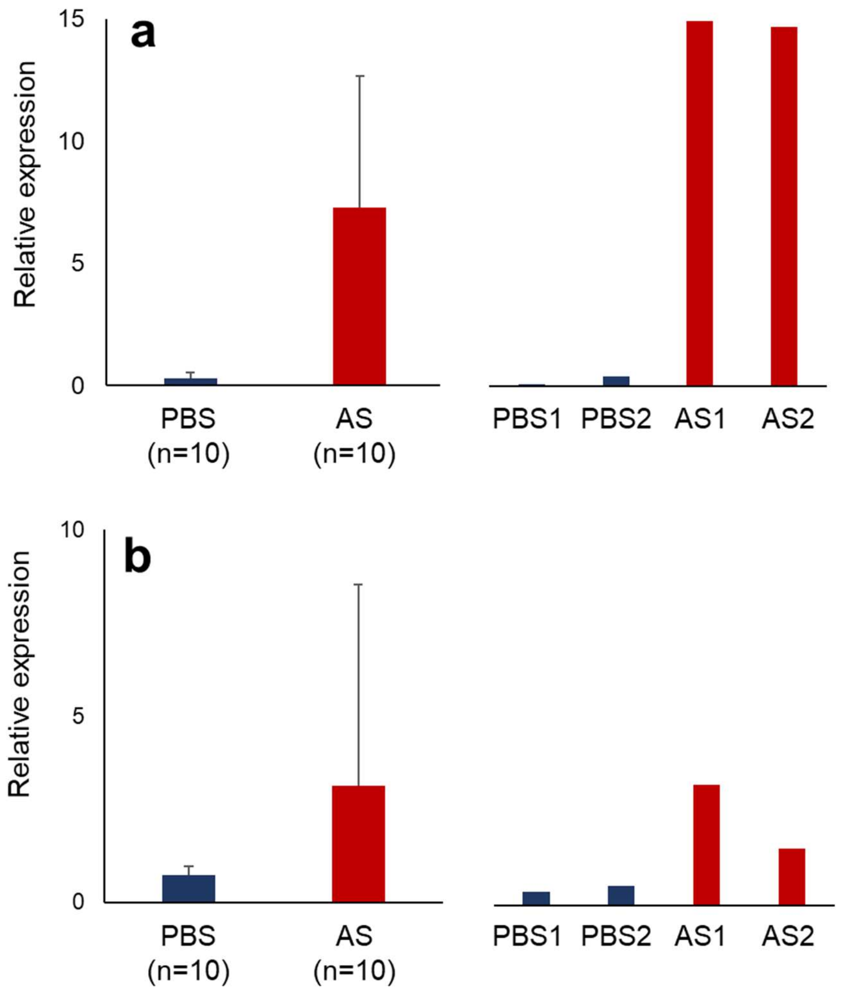


**Supplementary Figure 1**. Sample selection informed by expression of marker genes for the liver acute phase response. Quantitative PCR was used to measure the expression of established markers for the liver acute phase response in control (PBS injected) and *Aeromonas* infected (AS) samples: *saa* encoding serum amyloid alpha (**a**) and *hamp* encoding hepcidin (**b**). The data shown on the left of each figure is the average and standard deviation for normalized relative expression across the sampled population (n=10 per treatment). The data shown on the right shows normalized relative expression for the individual selected samples.

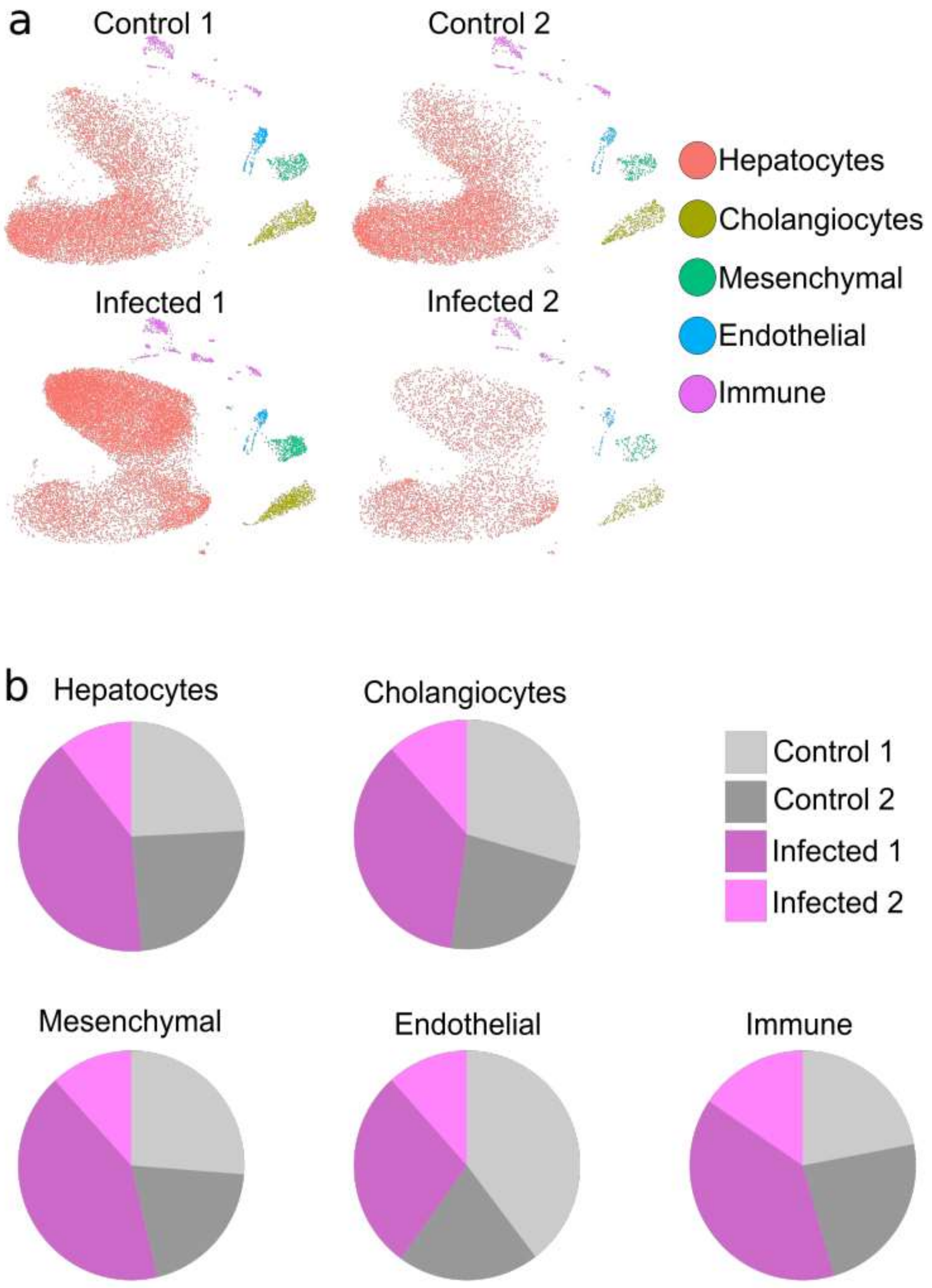


**Supplementary Figure 2**. Reproducibility of UMAP clustering across n=4 samples (**a**) and the contribution of each sample to the nuclei captured (**b**), for the five major liver cell clusters.

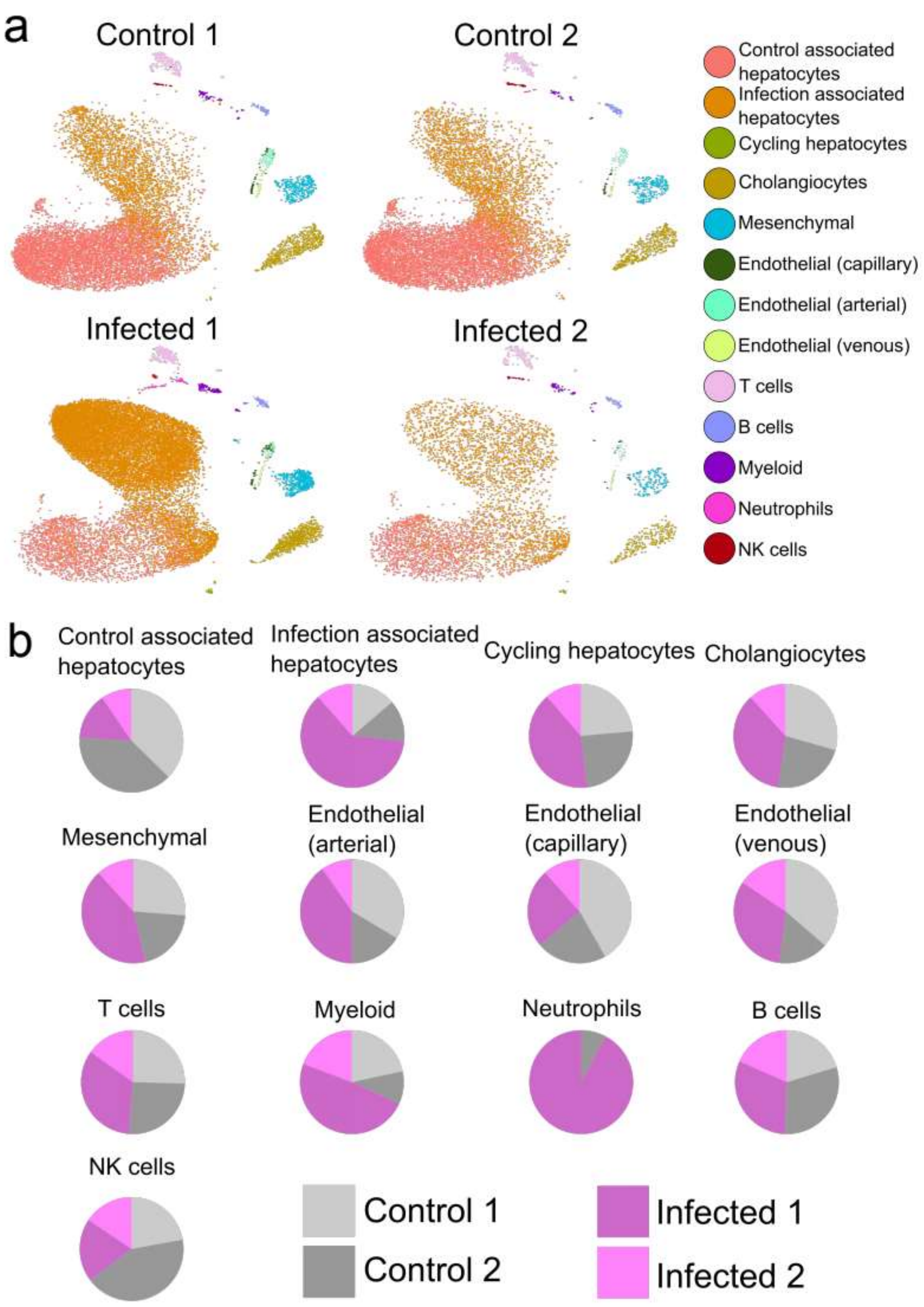


**Supplementary Figure 3**. Reproducibility of UMAP clustering across n=4 samples (**a**) and the contribution of each sample to the nuclei captured (**b**), for 13 cell clusters presented in Fig. 1a

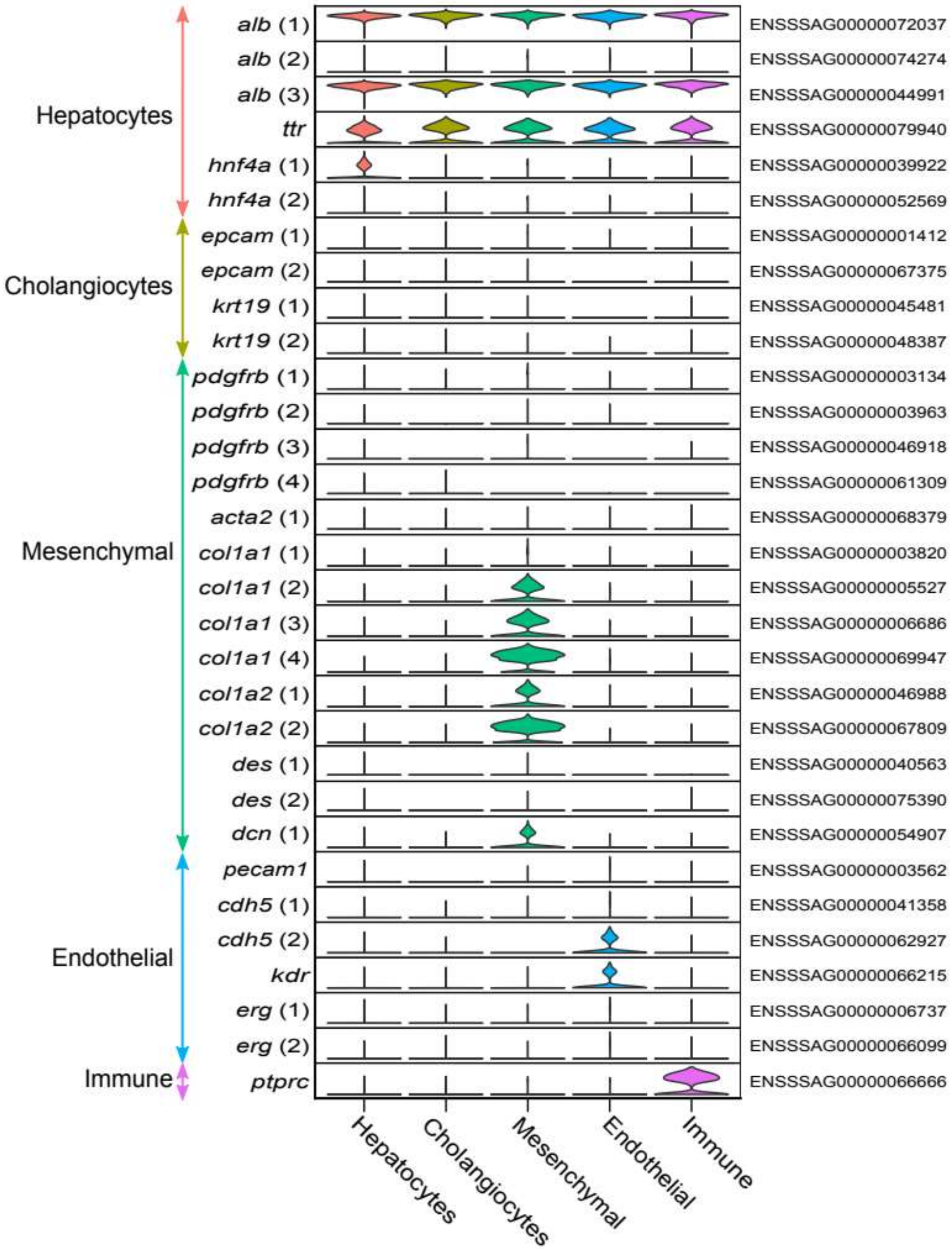


**Supplementary Figure 4**. Expression of Atlantic salmon orthologues to mammalian marker genes for the major liver cell types in our snRNA-Seq dataset. The genes shown are Atlantic salmon orthologues of marker genes for human liver cell types presented in Ramachandran et al. 2019 (cited in main text). In addition to the genes displayed, zero expression was observed for Atlantic salmon orthologues of the mesenchyme markers *Acta2* (ENSSSAG00000079015), *Dcn* (ENSSSAG00000061134), and *Des* (ENSSSAG00000047730 and ENSSSAG00000047730), in addition to the hepatocyte marker *Tf* (ENSSSAG00000080023). No orthologue of the hepatocyte marker *Cyp2a6*, mesenchyme marker *Col3a1*, and cholangiocyte marker *Cd24* could be identified in the Atlantic salmon genome.

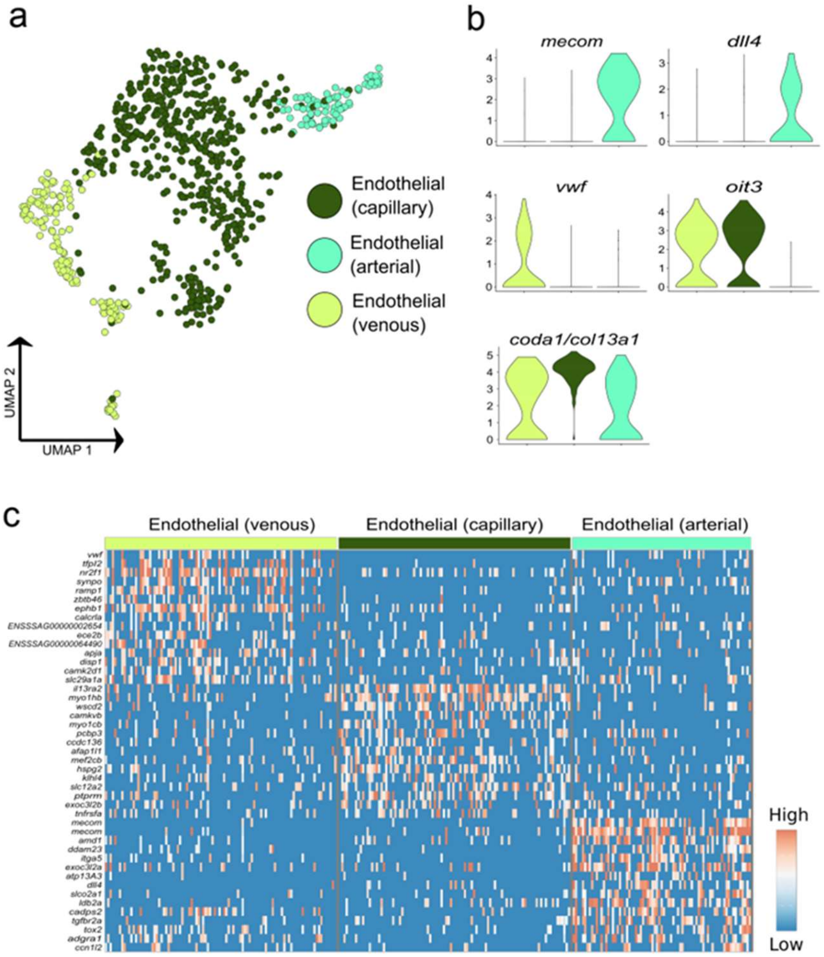


**Supplementary Figure 5.** Visualisation of the sub-clustering performed on the endothelial cells. (**a**) UMAP highlighting the three endothelial sub-clusters. (**b**) Expression of *mecom* (ENSSSAG00000041461) and *dll4* (ENSSSAG00000001347) indicate arterial properties; expression of *vwf* (ENSSSAG00000057945) indicates venous properties; expression of *oit3* (ENSSSAG00000054635) and *coda1* (ENSSSAG00000074170) indicate capillary characteristics. (**c**) Heatmap for the top 15 markers genes distinguishing each endothelial population based on a differential gene expression test.

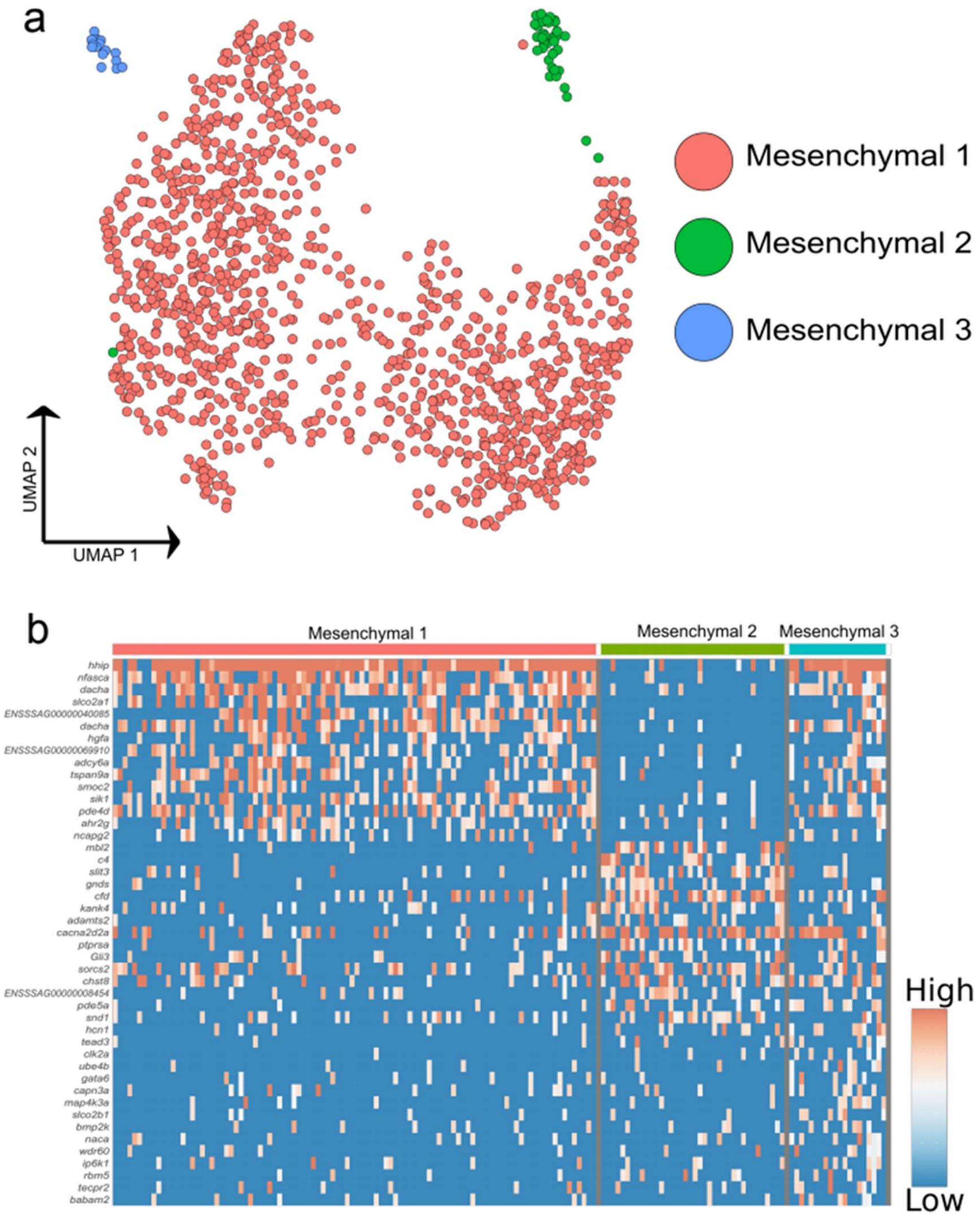


**Supplementary Figure 6.** Visualisation of the sub-clustering performed on the mesenchymal cells. (**a**) UMAP highlighting the three potential mesenchymal populations. (**b**) Heatmap for the top 15 markers genes distinguishing each mesenchymal sub-population.

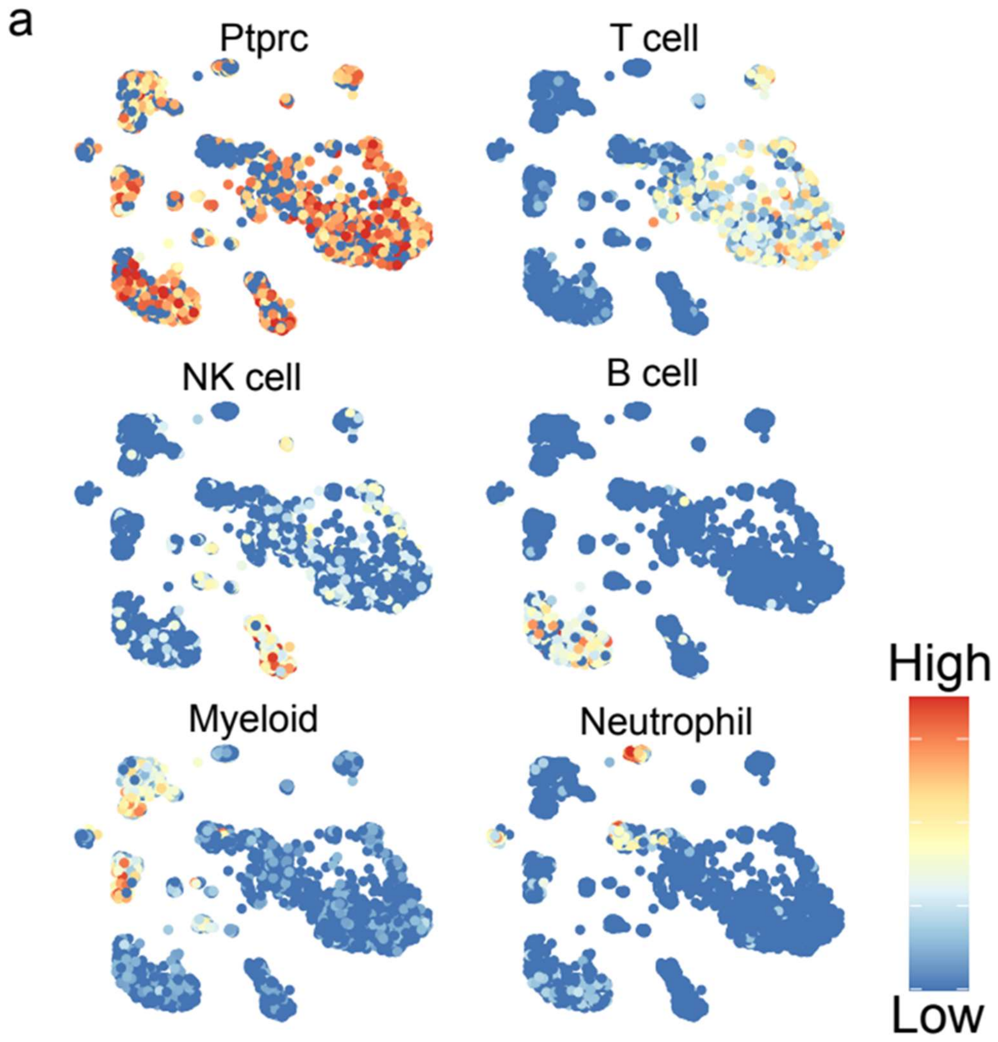


**Supplementary Figure 7.** Mean expression of *a priori* marker genes used to identify each broad immune cell population. Pan-immune marker *ptprc* was used to establish cells as immune cells, and mean expression levels of the following panel of markers was used to annotate with immune cell subtype. T cells: *tox2*, *tcf7*, *cd3e*; NK-like cell: *prf1.3* and *runx3*; B cells: *cd37* and *cd79a*; Myeloid cells: *mpeg1*, *cd63*, *csf1r*, and *lyz2*; Neutrophil: *ncf1*, *mmp9* and *itgax*.
